# Refined detection and phasing of structural aberrations in pediatric acute lymphoblastic leukemia by linked-read whole genome sequencing

**DOI:** 10.1101/375659

**Authors:** Jessica Nordlund, Yanara Marincevic-Zuniga, Lucia Cavelier, Amanda Raine, Tom Martin, Anders Lundmark, Jonas Abrahamsson, Ulrika Norén-Nyström, Gudmar Lönnerholm, Ann-Christine Syvänen

**Affiliations:** Department of Medical Sciences, Molecular Medicine and Science for Life Laboratory, Uppsala University, Sweden; Department of Immunology, Genetics and Pathology and Science for Life Laboratory, Uppsala University, Sweden; Department of Pediatrics, Institution for Clinical Sciences, Sahlgrenska Academy, Gothenburg University, Gothenburg, Sweden; Department of Clinical Sciences and Pediatrics, University of Umeå, Sweden; Department of Women’s and Children’s Health, Pediatric Oncology, Uppsala University, Sweden

**Keywords:** Childhood acute lymphoblastic leukemia, next generation sequencing, linked-read WGS, fusion gene, structural variants

## Abstract

Structural chromosomal rearrangements that may lead to in-frame gene-fusions represent a leading source of information for diagnosis, risk stratification, and prognosis in pediatric acute lymphoblastic leukemia (ALL). However, short-read whole genome sequencing (WGS) technologies struggle to accurately identify and phase such large-scale chromosomal aberrations in cancer genomes. We therefore evaluated linked-read WGS for detection of chromosomal rearrangements in an ALL cell line (REH) and primary samples of varying DNA quality from 12 patients diagnosed with ALL. We assessed the effect of input DNA quality on phased haplotype block size and the detectability of copy number aberrations (CNAs) and structural variants (SVs). Biobanked DNA isolated by standard column-based extraction methods was sufficient to detect chromosomal rearrangements even at low 10x sequencing coverage. Linked-read WGS enabled precise, allele-specific, digital karyotyping at a base-pair resolution for a wide range of structural variants including complex rearrangements and aneuploidy assessment. With use of haplotype information from the linked-reads, we also identified additional structural variants, such as a compound heterozygous deletion of *ERG* in a patient with the *DUX4-IGH* fusion gene. Thus, linked-read WGS allows detection of important pathogenic variants in ALL genomes at a resolution beyond that of traditional karyotyping or short-read WGS.

## INTRODUCTION

Our ability to sequence complete human genomes has increased owing to next generation sequencing (NGS) technologies, but detecting the whole spectrum of somatic single nucleotide variants (SNVs), copy number alterations (CNAs), and structural variations (SVs) in cancer cells is still challenging as SVs and CNAs remain the most difficult variant classes to discern in genomic data (1). A major reason for this limitation is that the human genome is diploid, consisting of a maternal and a paternal set of homologous chromosomes, and molecular haplotyping of alleles across large genomic regions is beyond the resolution of current short-read NGS technologies (2).

New “linked-read” technology, by which single molecules are massively barcoded in a microfluidic format and subsequently sequenced using short-read NGS technology, allows determination of molecular haplotypes across mega-base regions of the genome (3,4). An advantage of linked-read whole genome sequencing (WGS) over standard short-read WGS is its enhanced ability to detect the breakpoints of large-scale SVs and to provide long-range haplotype information for phasing. The long, linked-reads have enabled the assignment of complex structural variants and chromosomal rearrangements to individual chromosomes in germline and cancer genomes (3,5,6). Thus, linked-read WGS has the potential to overcome some of the limitations of short-read WGS for gaining a complete view of the structure of all genetic variants in a genome.

Structural chromosomal rearrangements that may lead to aberrant gene-fusions represent a leading source of information for diagnosis, risk stratification and prognosis in pediatric acute lymphoblastic leukemia (ALL) (7). Several chromosomal aberrations are recurrent in ALL and are used for classification of genetic subtypes associated with clinical outcome (8,9). The standard methods applied in clinical genetics laboratories today, such as karyotyping (G-banding) and fluorescent *in situ* hybridization (FISH), do not adequately capture the full spectrum of complex aberrations in the ALL genomes. Thus, up to 30% of B-cell precursor ALL (BCP-ALL) patients remain cytogenetically unclassified (10). WGS and whole-transcriptome sequencing (RNA-sequencing) technologies have enabled discovery of mutations, structural aberrations, and expressed gene-fusions in ALL (11–13). Recent large-scale RNA-sequencing studies have identified recurrent fusion genes with biological and clinical implications, such as those characterized by *DUX4, ZNF384*, and *MEF2D* rearrangements (14–19). However, only limited information is available on the chromosomal aberrations that are at the source of the gene-fusions and there are likely undetected structural aberrations with clinical importance yet to be discovered.

In the present study we evaluated if linked-read WGS technology could achieve the same level of detection as joint G-banding and FISH in a single linked-read WGS experiment. In our evaluation we focused on pathogenic structural aberrations in a set of well-characterized patients with pediatric ALL.

## MATERIALS AND METHODS

### Patient samples

This study included diagnostic samples from 12 children with acute lymphoblastic leukemia (ALL) enrolled on the Nordic Society of Pediatric Hematology and Oncology (NOPHO) protocols during 1998–2008 (8,20) and the t(12;21) cell line REH (21). Primary ALL samples were collected as described previously (22). The patients were selected from a large cohort of pediatric ALL patients based on presence of cytogenetic aberrations detected at diagnosis or expressed fusion genes detected by previous WGS or RNA-sequencing studies as well as availability of material from high blast count samples **(Supplementary Table S1) (11,18,19)**. DNA and RNA were extracted from 2–10 million cells using the AllPrep DNA/RNA Mini Kit (Qiagen) or the MagAttract HMW DNA kit (Qiagen). Fifty nanograms of DNA from ALL_370 was subjected to whole genome amplification with the DNA REPLI-g Midi Kit (Qiagen). The DNA concentrations were measured using the Qubit dsDNA Broad Range assay (Invitrogen). The study was approved by the Regional Ethics Review Board in Uppsala, Sweden and was conducted according to the guidelines of the Declaration of Helsinki. The patients or their guardians provided informed consent.

### Karyotyping and molecular diagnosis

ALL diagnosis was established by analysis of leukemic cells with respect to morphology, immunophenotype, and cytogenetic aberrations. High hyperdiploidy (HeH) was defined as presence of 51–67 chromosomes per cell (23). FISH or RT-PCR analyses were used to screen for t(12;21)(p13;q22)[*ETV6-RUNX1*] and t(9;22)(q34;q11)[*BCR-ABL1*]. Whole chromosome paint (Metasystems XCP orange/green XCyting Chromosome Paints) and subtelomeric probes (Vysis Totelvysion probes) were used to validate translocations on metaphase spreads from cultured bone marrow cells from patients ALL_559, ALL_707 and ALL_386. The hybridized slides were analyzed using a Zeiss fluorescence microscope (Carl Zeiss) and chromosome-colored images were captured using the Isis software (MetaSystems).

### Library construction and sequencing

Sequencing libraries were prepared from 1-1.2 ng of genomic DNA according to the manufacturer’s instructions for preparation of GemCode and Chromium WGS libraries (10x Genomics). The DNA molecules were partitioned into droplets including a GemCode/Chromium barcode-specific gel bead (GEM) and subsequently amplified by PCR with combined adaptor tagging using GemCode/Chromium barcodes. The droplets were fractured to release the barcoded PCR products and subjected to construction of indexed libraries. GemCode libraries were sequenced on an Illumina HiSeq2500 instrument with a customized sequencing protocol (read1:98bp, i7:8bp, i5:14bp, read2:98) to an average depth of 14x. Chromium libraries were sequenced on an Illumina HiSeqX instrument with 150 bp paired-end reads to an average depth of 32x.

### Linked-read data analysis

Linked-read WGS data was processed and analyzed using the Long Ranger pipeline from 10x Genomics (v1.2.0 for GemCode and v2.1.6 for Chromium) with the hg19/GRCh37 reference genome. The ‘-somatic’ flag was used for the Chromium libraries. Data were visualized using the Loupe Genome Browser v2.1.1. Phasing haplotype reconstruction was performed by Long Ranger. SVs called by Long Ranger were manually reviewed and assessed against karyotype data, CNA data from Illumina Infinium arrays, and fusion genes detected by RNA-sequencing. Genomic copy number (CN) levels were estimated by chromosomal segmentation read-depth analysis in 10 Kb windows using the CNVnator software (24). B-allele frequencies were calculated from VCF files from Long Ranger using the VariantAnnotation package and custom scripts in R (25). Ideograms of derived chromosomes were drawn to scale with the CyDAS software (26).

### RNA-sequencing

RNA-sequencing libraries from REH and ALL_402 were constructed from 300ng total RNA with the TruSeq stranded total RNA protocol (RiboZero human/mouse/rat) according to the manufacturer’s instructions (Illumina). The libraries were sequenced on a NovaSeq 6000 instrument with 100 bp paired-end reads. Strand-specific RNA-sequencing data was previously been generated for all the remaining patient samples in the study, except from patient ALL_370 (11,18,19). Fusion genes were called and validated using a previously descried approach (19) based on FusionCatcher 0.99.7d (27).

### Copy Number Analysis

Previously generated data from Infinium HumanMethylation450 BeadChips (450k arrays) are available at the Gene Expression Omnibus (GSE49031) (28). The R package “CopyNumber450kCancer” was used to detect CN alterations in the 450k array data (29). Genomic DNA (200ng) from nine patient samples was subjected to genotyping on the Illumina HumanOmni2.5 Exome-8v1 SNP arrays according to the manufacturer’s specifications (Illumina). CN alterations were called from the SNP array data using the Tumor Aberration Prediction Suite (30).

## RESULTS

Eighteen sequencing libraries of which 13 were prepared with GemCode reagents and five were prepared with Chromium reagents were sequenced from 12 primary samples collected from pediatric ALL patients at diagnosis and from the REH ALL cell line using different types of input DNA (**Table 1**). The 18 libraries were sequenced to an average coverage of 14x and 32x for GemCode and Chromium, respectively. The number of phased SNPs ranged from 69-99% (mean 93%) and the longest phase blocks ranged from 0.3-18 megabases (Mb) in size (mean size 7 Mb) (**Supplementary Table S2**). The quality and type of input DNA as well as the sequencing coverage had the largest effect on the length and quality of phasing, however the majority of large structural aberrations in the ALL genomes remained detectable in un-phased regions in lower coverage libraries.

**Table 1.** Patient characteristics and linked-read WGS library statistics.

| Patient ID | Subtype | Library Name | DNA extraction method | Library Type | Average sequencing depth | Weighted mean molecule length (Kb) | SNPs phased | Longest phase block (Mb) | N50 phase block (Mb) | Large structural variant calls | Revised karyotype after linked-read WGS<sup>a</sup> |
| --- | --- | --- | --- | --- | --- | --- | --- | --- | --- | --- | --- |
| ALL_370 | HeH | ALL_370.1 | Standard column + WGA | GemCode | 18 | 28 | 0.69 | 0.28 | 0.01 | 0 | 55,XX,+X,+4,+6,+10,+14,+17,+18,+21,+21 |
|  |  | ALL_370.2 | Standard column | GemCode | 21 | 49 | 0.83 | 0.92 | 0.06 | 2 |  |
| ALL_689 | HeH | ALL_689 | Standard column | GemCode | 22 | 70 | 0.97 | 9.87 | 1.37 | 11 | **54,XX,+X,dup(1)(q24q42),+4,+6,+10,+14,+17,+18,+21** |
| ALL_47 | HeH | ALL_47.2 | Standard column | Chromium | 33 | 60 | 0.99 | 14.15 | 2.27 | 30 | **<u>58,XY,+4,+5,+6,+9,+10,+12,+14,+17,+18,-19,+19,+21,+21,+22</u>** |
|  |  | ALL_47.1 | Standard column | GemCode | 21 | 66 | 0.96 | 7.26 | 1.26 | 9 |  |
| ALL_458 | ETV6-RUNX1 | ALL_458 | Standard column | GemCode | 9 | 56 | 0.82 | 3.36 | 0.44 | 5 | **<u>47,XY,+10,del(11)(q22.1q25),t(12;21)(p13.2;q22),del(12)(p12.1p13.2)</u>** |
| ALL_386 | ETV6-RUNX1 | ALL_386 | Standard column | Chromium | 26 | 37 | 0.98 | 5.00 | 0.80 | 424 | **<u>46,XY,del(2)(q33.1q37.3),der(3)del(3)(p21.2p21.31)t(3;12)(p21.31;q24)ins(3;3)(q21.2;p21.31p21.31),der(12)t(14;12)(q24.1;p13.2)t(3;12)(q21.3;q24.11),del(12)(p13.2),der(14)t(14;2)(q24.1;q37.3)t(2;21)(q33.1;q22.12),del(19)(q13.32q13.43),der(21)t(12;21)(p13.2;q22.12)</u>** |
| REH | ETV6-RUNX1 | REH.1 | MagAttract - HMW | Gemcode | 11 | 128 | 0.90 | 14.02 | 2.24 | 29 | **<u>46,X,-X,del(3)(p14.2p22.3),der(4)t(4;16)(q32.1;q24.3),t(5;12)(q23.2;p13.2),der(12)inv(12)(p12q24)t(4;12)(q32;q23),der(16)t(16;21)(q24.3;q22.12),+16,der(21)t(12;21)(p13.2;q22.12)</u>** |
|  |  | REH.2 | Standard column | Gemcode | 10 | 62 | 0.92 | 7.58 | 1.14 | 24 |  |
| ALL_402 | BCR-ABL1 | ALL_402.1 | MagAttract - HMW | Gemcode | 11 | 126 | 0.92 | 18.04 | 3.27 | 12 | 46,XY,t(1;5;9;22)(**<u>p36.33;q31.2;q34.12;q11.23</u>**),**<u>del(8)(p11p23),dup(8)(p11.23q24.3)</u>**,del(9)(p21p21) |
|  |  | ALL_402.2 | Standard column | Gemcode | 10 | 42 | 0.84 | 2.03 | 0.28 | 6 |  |
| ALL_390 | DUX4-IGH | ALL_390.1 | Standard column | GemCode | 21 | 62 | 0.97 | 7.65 | 1.27 | 6 | 46,XX,**<u>del(6)(q14.1q27)</u>** |
|  |  | ALL_390.2 | Standard column | Chromium | 31 | 50 | 0.99 | 9.46 | 1.87 | 32 |  |
| ALL_501 | DUX4-IGH | ALL_501 | Standard column | GemCode | 10 | 59 | 0.89 | 3.50 | 0.58 | 6 | 46,XX |
| ALL_604 | TCF3-ZNF384 | ALL_604 | Standard column | Chromium | 32 | 49 | 0.99 | 10.74 | 1.76 | 46 | 46,XY,del(6)(**<u>q16.2q22.33</u>**),del(7)(**<u>q21.3q36.3</u>**),t(12;19)(p13.31;p13.3) |
| ALL_613 | EP300-ZNF384 | ALL_613 | Standard column | Chromium | 36 | 52 | 0.99 | 11.47 | 1.85 | 48 | 46,XY,**<u>dup(1)(q21q44),t(12;22)(p13.2;q13.2)</u>**,del(16)(q21q24.3) |
| ALL_707 | PAX5-ELN | ALL_707 | Standard column | GemCode | 10 | 60 | 0.88 | 5.23 | 0.63 | 5 | 46,XY,**<u>del(7)(q11)</u>**,der(9)t(7;9)(q11;p13),del(9)(p13p24) |
| ALL_559 | T-ALL | ALL_559 | Standard column | GemCode | 9 | 61 | 0.81 | 2.36 | 0.43 | 8 | 46,XY,**<u>der(7)t(7;9)(q34;q31),t(7;11)(q34;p15),der(9)t(7;9)(q34;q31)</u>**del(9)(p21p21),del(9)(p21p21) |

<sup>a</sup> The parts of the karyotype revised after linked-read WGS are highlighted in bold and underlined.

### Effect of input DNA

High molecular weight DNA (HMW DNA) resulted in the longest phase blocks that spanned 14-18 Mb of DNA, whilst standard column-based DNA preparations that had undergone repeated freeze-thawing without selection for HMW DNA fragments yielded shorter phase blocks ranging from 1-14 Mb (**Table 1**). The sizes of the longest and the N50 phase blocks correlated with the average size of the input DNA estimated by the Long Ranger software (**Figure 1A-B**). Shorter haplotype blocks were generated for the libraries prepared by GemCode (average ~5 Mb) compared to libraries prepared using Chromium reagents (average ~10 Mb), likely due to lower sequencing depth of the Gemcode libraries.

**Figure 1.**
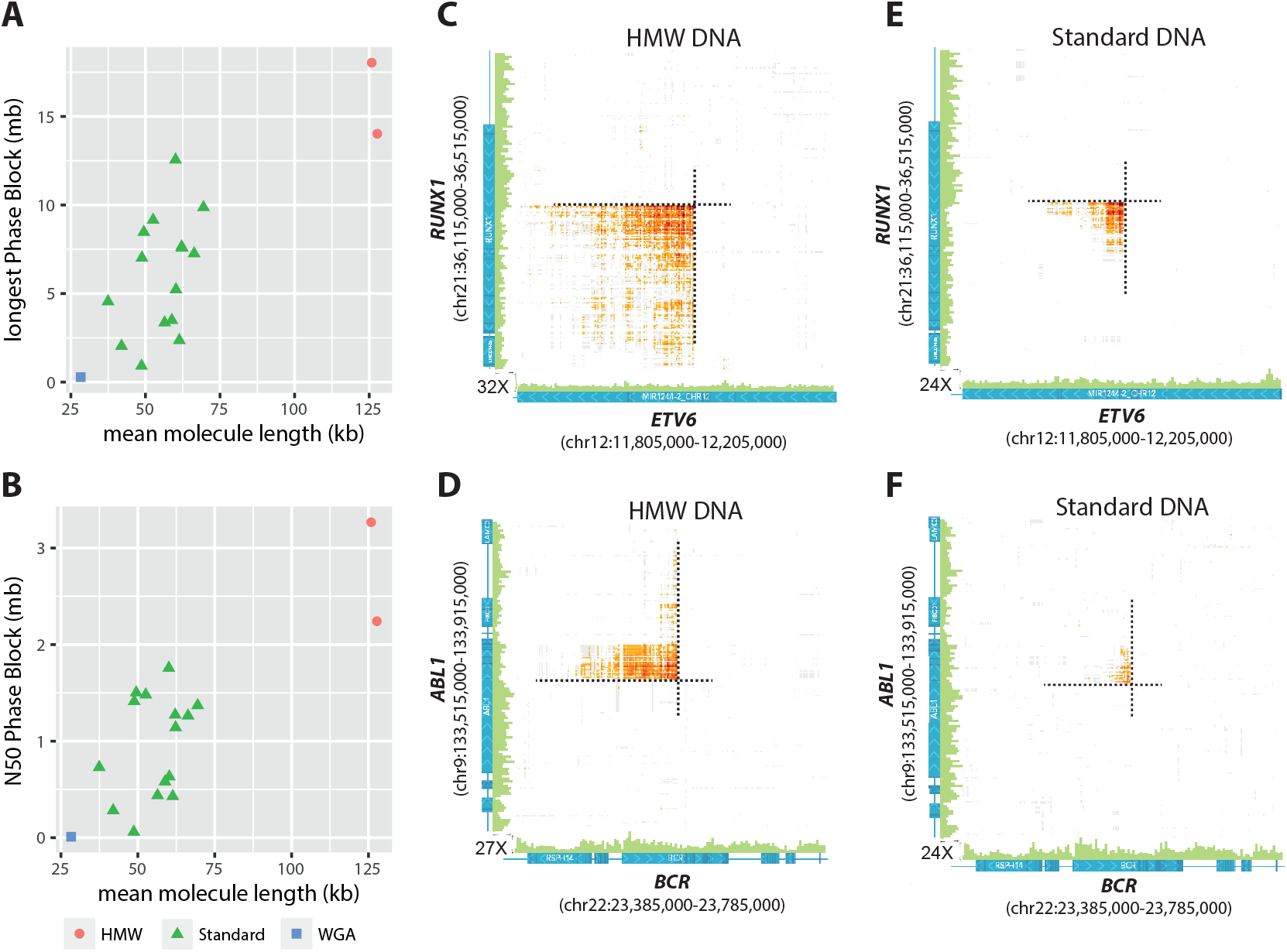
Effect of genomic DNA fragment size on phasing and detection of fusion genes. The weighted average DNA size estimated by the 10x Genomics Long Ranger software is plotted against (A) the size of the largest phase block and (B) the median size (N50) of the phase block. High molecular weight (HMW) DNA (red dots), DNA extracted with a column-based method (standard, green triangles), and DNA that was been whole-genome amplified (WGA, blue squares) are indicated in the plots. (C-F) Heatmaps of overlapping linked reads supporting inter-chromosomal translocations are plotted in orange (10x Genomics Loupe software). HMW DNA extracted from the (C) REH cell line harboring t(12;21) and patient ALL_402 harboring t(9;22) (D). Standard column extracted DNA from the REH cell line (E) and from patient ALL_402 (F). The expected breakpoints in the fusion genes were identified in both HMW standard column DNA extractions.

To directly compare the HMW and standard column-based DNA extractions, HMW DNA was freshly prepared from two samples harboring t(12;21) or t(9;22) and were compared to DNA extracted with a column-based method. GemCode libraries were prepared and sequenced to 10x coverage from each DNA sample. The expected genomic breakpoints were identified in both the HMW and standard DNA libraries (**Figure 1A-B**). However, the cumulative number of common barcodes shared between the two loci of each translocation yielded stronger signal intensities in the HMW DNA (60,044 and 26,316, **Figure 1C-D**, respectively) compared to the standard column-based DNA samples (12,733 and 7,500, **Figure 1E-F**, respectively).

To determine the utility of whole genome amplified (WGA) DNA for long-range phasing we compared the phasing performance of DNA that had been subjected to WGA (ALL_370.1) to that of standard genomic DNA (ALL_370.2). Not unexpectedly, the sample that had been subjected to WGA prior to sequencing library preparation (ALL_370.1) yielded the poorest results, with the longest haplotype block only 0.3 Mb in length and with only 69% of the SNPs phased (**Table 1**). Due to the poor performance of WGA DNA, we excluded the results from the WGA library ALL_370.1 from further analyses.

### Detection of chromosomal aberrations and copy number using linked-read WGS

For seven of the 13 individual ALL genomes analyzed, detailed karyotype information was available by G-banding or FISH for subtype-defining genetic aberrations such as high hyperdiploidy (HeH), t(12;21), or t(9;22). Thus we were able to verify the results from the linked-read WGS data by comparison to that obtained from genetic analysis at diagnosis. The somatic genomic variants in the remaining six patients with either T-ALL or B-other subtype were determined in previous studies by a combination of diagnostic karyotyping, WGS, RNA-sequencing, or Infinium DNA methylation/SNP arrays (11,19). These six patients had complex or incomplete karyotype information as detected by G-banding and FISH at ALL diagnosis, and thus we aimed to better resolve the aberrations in these ALL genomes. We used the Loupe software provided by 10x Genomics to screen for and identify the exact chromosomal breakpoints at base-pair resolution and the haplotypes of the large SVs in the ALL genomes. In all cases, we validated our findings using existing karyotype information, FISH (when cells were available), and/or a combination of Infinium arrays for copy number estimates and RNA-sequencing for detection of expressed fusion. The results for each patient and subtype are detailed below and in each case a revised karyotype after linked-read WGS is given in **Table 1**.

### High Hyperdiploidy (HeH)

Two patients (ALL_370 and ALL_689) included in our study had the classical HeH subtype with 55 chromosomes and no detectable translocations by karyotyping and/or FISH. The aneuploidy estimates in the karyotype were consistent with Infinium array data for both patients (**Table 1**). Using the linked-read WGS data, we binned the average sequencing coverage in 10 Kb bins across the genome and scanned for copy-number alterations (CNAs) (**Figure 2A-B**). The linked-read WGS estimates of copy numbers correlated perfectly with that from the arrays and with the karyotype for ALL_370 and ALL_689. A third patient (ALL_47) with “normal” (failed) karyotype at diagnosis was suspected to have the HeH subtype based on a previous study (18). For this patient we verified the HeH karyotype in the linked-read WGS data to be +4,+5,+6,+9,+10,+12,+14,+17,+18,-19,+19,+21,+21,+22, which was verified by CNA analysis (**Table 1; Figure 2C**). The copy neutral loss of chromosome 19 (uniparental disomy) was visible in the linked-read WGS data by an overrepresentation of homozygous SNVs on chromosome 19 (**Supplementary Figure S1**). The chromosomal copy number alterations were successfully defined by linked-read WGS in each of the three HeH cases, with directly comparable results to traditional karyotyping and microarrays.

**Figure 2.**
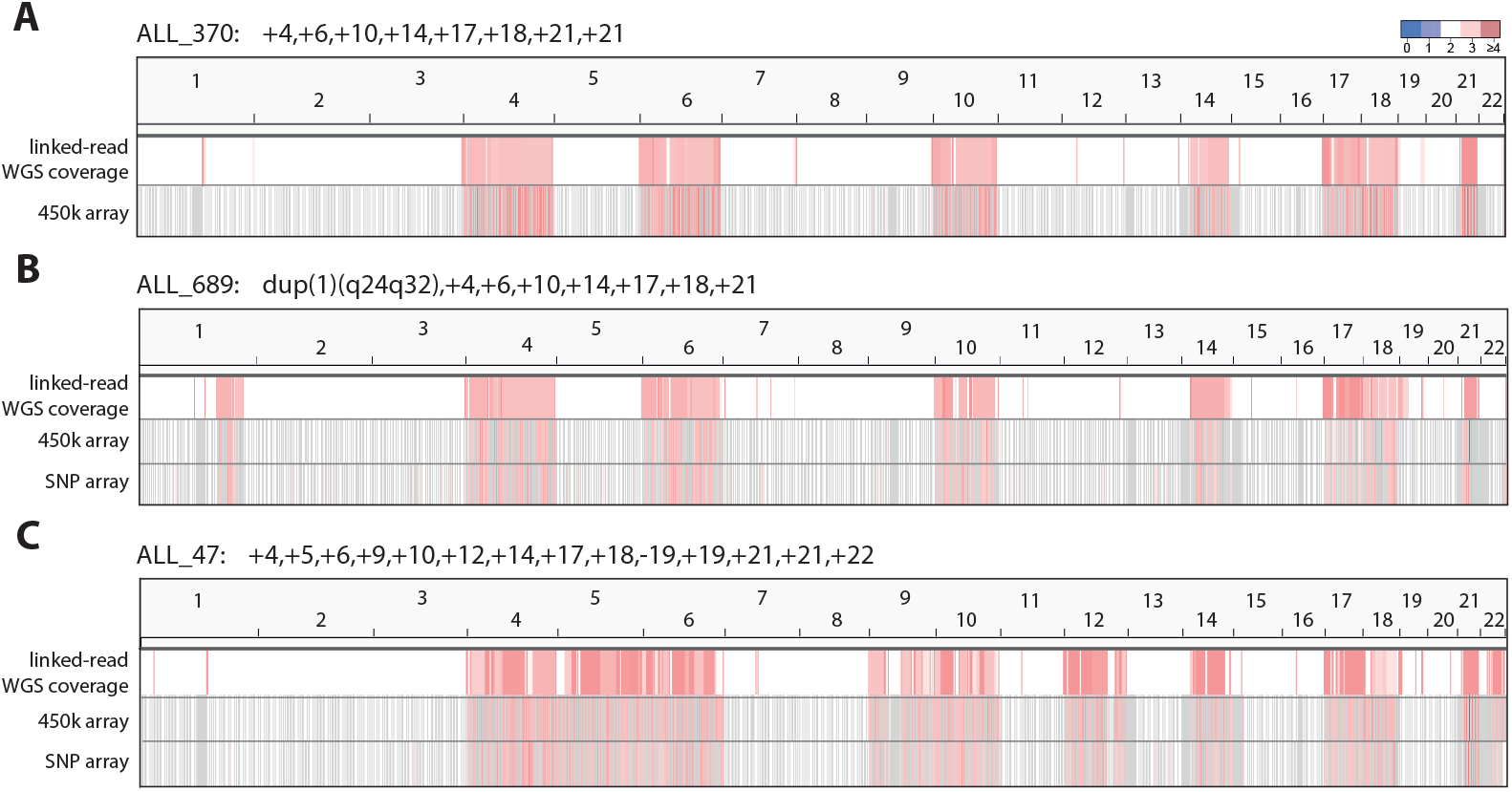
Copy number by chromosome for the three ALL patients with the HeH subtype (A-C). The average linked-read WGS coverage data calculated in 10kb bins is plotted in the top row of each panel. The Log R ratios from Infinium SNP and/or 450k array data are visualized in the lower part of each panel. Red coloring indicates chromosomal gains according to the color key above panel A.

### t(12;21) and t(9;22)

The t(12;21) translocation and associated aberrations were analyzed in two patients (ALL_386 and ALL_458) and the REH cell line (**Figure 3A-C**). As was anticipated from diagnostic karyotyping and previous short-read WGS of patient ALL_458 (11), a balanced t(12;21) translocation resulting in the expression of both the canonical *ETV6-RUNX1* and the reciprocal *RUNX1-ETV6* fusion genes was unambiguously detected at base-pair resolution in the linked-read WGS data (**Supplementary Figure S2**). A deletion spanning over a 2.1 Mb region that includes the second allele of *ETV6* was observed on the other haplotype. Besides gain of chromosome 10 and a 38 Mb deletion of chromosome 11q22-q25, no other large structural variants were identified in ALL_458.

**Figure 3.**
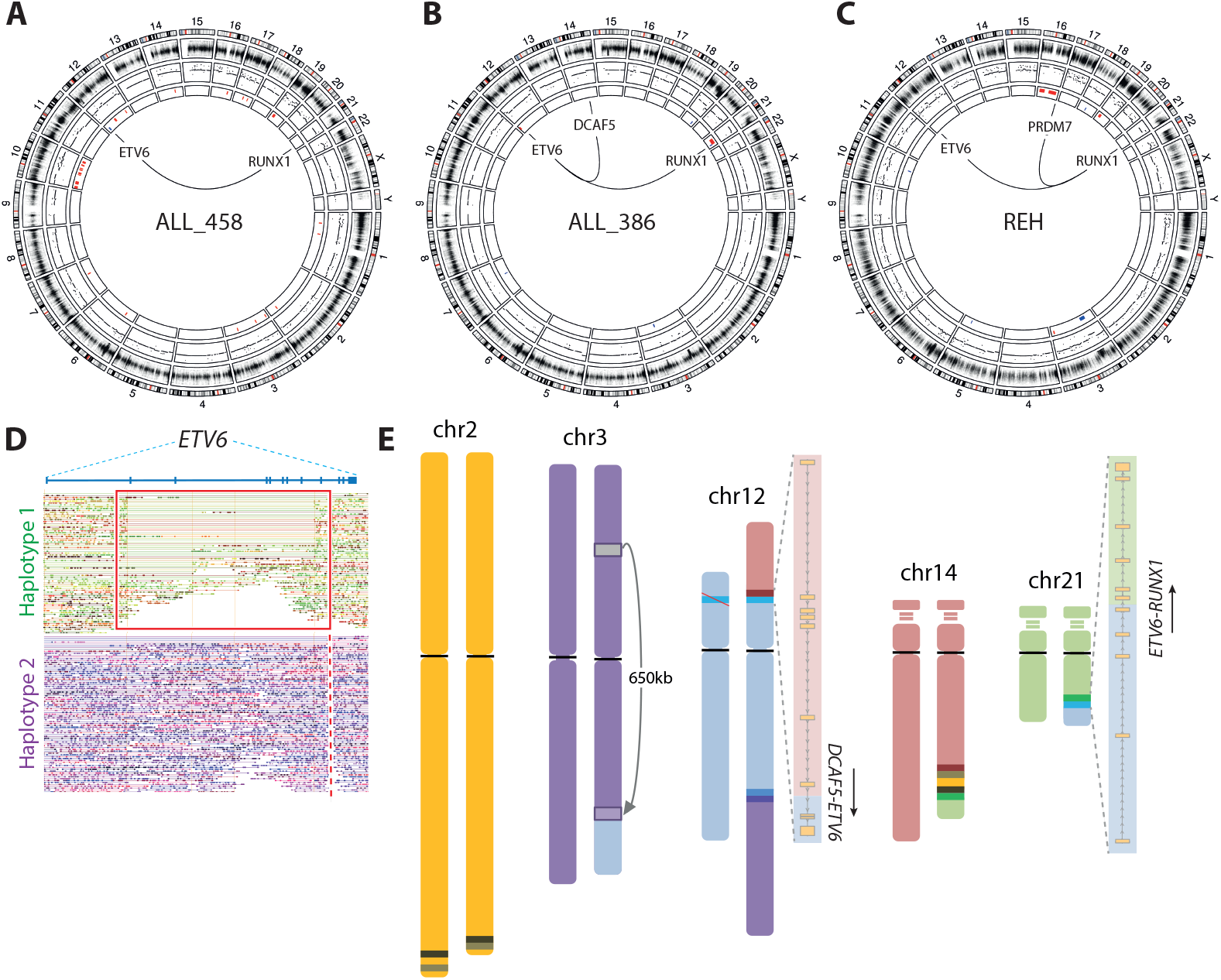
Structural aberrations detected by linked-read WGS in t(12;21)*ETV6-RUNX1* genomes. A-C) Circos plots for patients ALL_386, ALL_458 and the REH cell line. The first (outer) track shows the chromosomes and their banding, the second track shows log R ratios from Infinium arrays, the third track shows copy number determined by linked-read WGS in 10kb bins, and the fourth (innermost) track shows copy number calls using the CNVnator software. Red indicates amplifications and blue indicates deletions. Expressed fusion genes are highlighted within each circos plot, solid lines indicate in-frame fusion genes. D) Linked-reads mapped to two haplotypes at the *ETV6* locus in patient ALL_386, which depicts a deletion on haplotype 1 (indicated by the red box) and the breakpoint giving rise to the *DCAF5-ETV6* and the *ETV6-RUNX1* fusion genes is indicated by a dashed line on the other allele (haplotype 2). E) Schematic representation of the chromosomal rearrangements resulting in derived chromosomes as determined by linked-read WGS in ALL_386. The ideograms are drawn to scale using the CyDAS software. The resulting fusion transcripts with breakpoints are drawn alongside the chromosomes involved in the translocations.

In contrast, for ALL_386 and the REH cell line, karyotype data suggested complex re-arrangements involving *ETV6, RUNX1* and several other chromosomes. The diagnostic karyotype for patient ALL_386 suggested a complex series of translocations including chromosomes 3, 12, 14 and 21, of which two of the translocations resulted in in-frame fusion genes: *DCAF5-ETV6* from a translocation between chromosomes 14q24.1 and 12p13.2 and *ETV6-RUNX1* from a translocation between 12p13.2 and 21q22.12 (19). The haplotype phasing information resolved an intragenic 0.15 Mb deletion of one allele of *ETV6* (haplotype 1) and that the two fusion genes (*ETV6-RUNX1* and *DCAF5-ETV6*) originated from the second allele (haplotype 2) of *ETV6* (**Figure 3D**). Linked-read WGS resolved the exact breakpoints on chromosomes 3, 12 14 and 21 that were expected, and identified several additional alterations that were missed by genetic analysis at diagnosis. Of which, *DCAF5* (chr14) and the reciprocal *RUNX1* (chr21) loci were separated by a 44 Mb insertion of a region originating from chromosome 2q on the derived chromosome 14q (**Figure 3E**). Furthermore, a 650 kb region from chromosome 3p21.31 was inverted and inserted into the derived chromosome 3q21.2 arm where the material from chromosome 12p was translocated. A schematic overview of the derived chromosomes determined by linked-read WGS and validated by FISH is outlined in **Supplementary Figures S3-S4**.

The structure of the complex rearrangement involving chromosomes 4, 5, 12, 16 and 21 in the REH cell line as determined by linked-read WGS was consistent with the reported karyotype (**Supplementary Figure S5**). Both alleles of chromosomes 12 were involved in translocations with breakpoints occurring in proximity of, or within, the *ETV6* gene. One allele of chromosome 12 is involved in a balanced t(5;12) translocation with a flanking 0.6 Mb deletion at the *ETV6* locus. The second allele was involved in a series of translocations between chromosomes 4, 12, 21, and 16 in which the *ETV6-RUNX1* fusion gene was formed. RNA-sequencing confirmed expression of *ETV6-RUNX1* from t(12;21) and *RUNX1-PRDM7* from t(16;21), but no expressed fusion genes were observed arising from the other breakpoints (**Figure 3C**). The inversion of chromosome 12p13.2-q23.1 expected from the karyotype was observed in the linked-read data. A duplication of chromosome 18 and an inversion of chromosome 5 were expected according to the REH karyotype, however we did not find evidence of these aberrations in the linked-read WGS or Infinium array data.

The genome of patient ALL_402 was expected to harbor the t(9;22) translocation. Unexpectedly, linked-read WGS revealed a much more complex rearrangement involving the *BCR* (22q11.23), *ABL1* (9q34.12), *PRRC2B* (9q34.13), *SIL1* (5q31.2) and *LINC01128* (1p36.33) loci (**Supplementary Figure S6**). In addition to the deletion of chromosome 9p21 reported in the karyotype, on chromosome 8 we detected a 35 Mb deletion (8p11.23-p23.3) and an amplification starting at 8p11.23 and continuing through the entire q-arm (**Figure 4A**). RNA-sequencing verified that the 5’ end of *BCR* is fused with the 3’ end of *ABL1*, the 5’ ends of the reciprocal *ABL1* and *SIL1* loci form a head to head translocation resulting in two truncated transcripts, the 5’ end of *LINC01128* is fused with the 3’ end of *SIL1*, whilst the 5’ end of *PRRC2B* is fused with the reciprocal 3’ end of the *BCR* gene (**Figure 4B**). None of the complex rearrangements were phased in the linked-read sequencing data, but in this instance the phasing information was not necessary to fully resolve the structure of the breakpoints.

**Figure 4.**
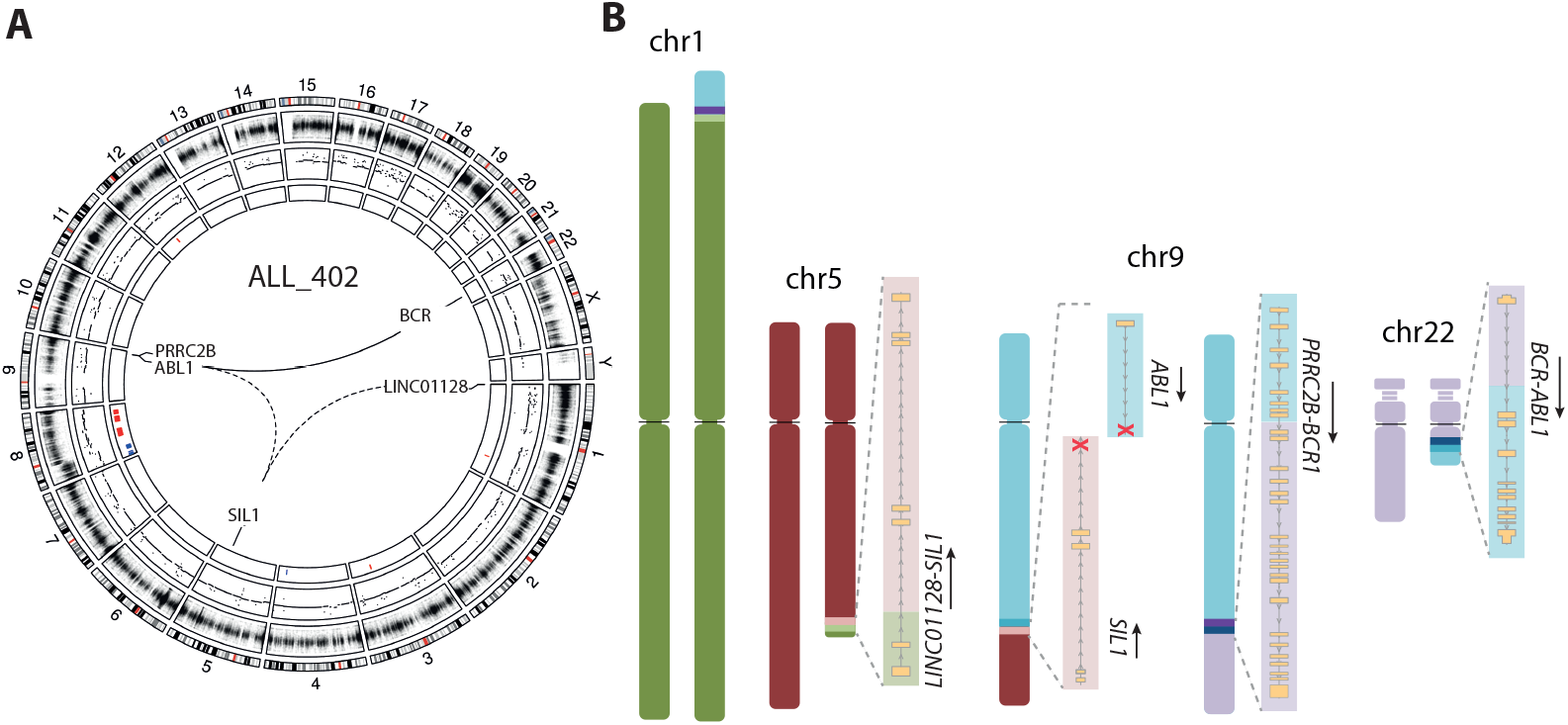
Complex structural rearrangements in the patient ALL_402. A) A circos plot depicting the genome-wide copy number changes in ALL_402. The first (outer) track shows each chromosome and their banding, the second track shows log R ratios from infinium arrays, the third track shows copy number determined by linked-read WGS in 10kb bins, and the fourth (innermost) track shows copy number calls using the CNVnator software. Red indicates amplifications and blue indicates deletions. Expressed fusion genes are highlighted inside of the circos plot, solid lines indicate in-frame and dashed lines indicate out of frame fusion or truncated genes. B) The derived chromosomes as outlined using linked-read WGS. The structures of the expressed fusion genes are shown alongside their derived chromosomes with the direction of transcription indicated by arrows. The ideograms are drawn to scale using the CyDAS software.

Taken together, we show that linked-read WGS has the power to detect both aneuploidies and translocations such as t(12;21) and t(9;22), and highly complex translocations that were either missed or miss-annotated by traditional karyotype analysis by G-banding and FISH at ALL diagnosis.

### B-other

*DUX4* and *ZNF384*-rearrangements define newly described subtypes of BCP-ALL that were initially detected in large-scale RNA-sequencing studies (15,16,31). The *DUX4-IGH* fusion gene results from an insertion of the *DUX4* gene (subtelomeric region of chr4q and 10q), into the enhancer region of the *IGH* locus (chr14) (32). Prior to discovery of the *DUX4* rearrangement, this putative subgroup was referred to as “*ERG*-deleted” because of a high prevalence of *ERG* deletions and an otherwise normal karyotype (33,34). As was expected for the two patients with *DUX4-IGH* (ALL_390 and ALL_501), largely normal karyotypes were observed, with the exception of a large 93 Mb deletion on chromosome 6q14.1-q27 in ALL_390 (**Supplementary Figure S7**). A large 6.5 Mb phase block on chromosome 21q22 enabled detection of a compound heterozygous deletion of *ERG* transcript variant 1 (NCBI Reference Sequence: NM_182918.3) in ALL_390 with a 57.2 Kb deletion on haplotype 2 spanning exons 3-10 and a second 9.3 Kb focal deletion of exon 1 on haplotype 1 (**Figure 5A**). The Long Ranger software was not able to resolve the insertion of the 1.2 Kb *DUX4* gene into the enhancer region of the *IGH* locus in either patient. However, upon visual examination of raw reads with the aid of the Integrated Genome Viewer, we were able to identify split linked-reads supporting the insertion of at least one copy of *DUX4* into the *IGH* locus in the Chromium library ALL_390.2, but not in the lower coverage GemCode libraries ALL_390.1 or ALL_501 (**Supplementary Figure S8**).

**Figure 5.**
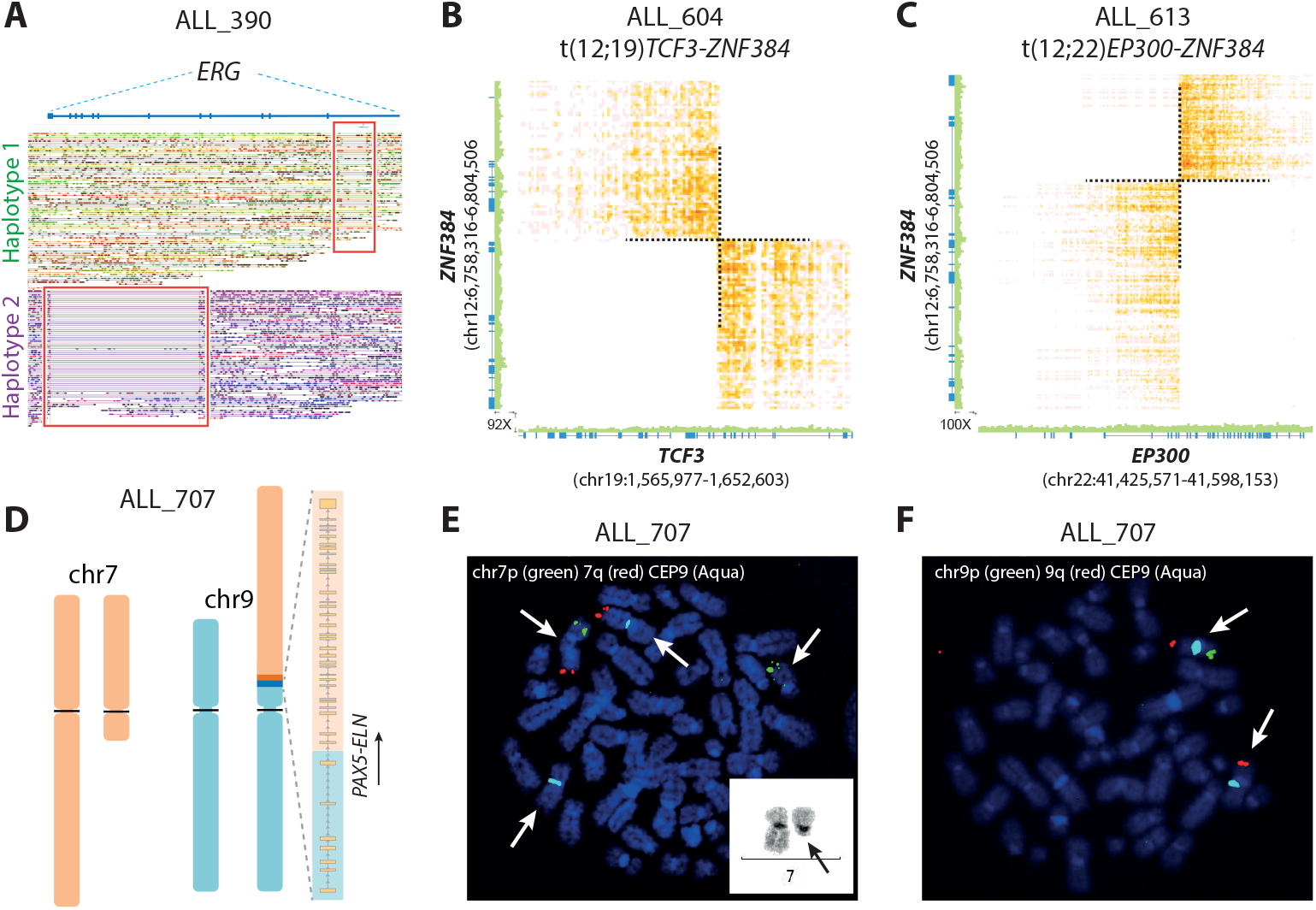
Structural rearrangements detected in B-other patients by linked-read WGS. A) Linked-reads mapped to each of the two homologous chromosomes at the *ERG* locus on chromosome 21 in patient ALL_390. Reads are color-coded by chromosome and deletions are marked by red squares. B-C) Heatmaps of overlapping linked-reads supporting subtype-defining balanced inter-chromosomal translocations from the 10x Genomic’s Loupe software. (B) The genomic breakpoint in chromosomes 12 and 19, resulting in the *TCF3-ZNF384* fusion gene in patient ALL_604. (C) The genomic breakpoint in chromosomes 12 and 22, resulting in the *EP300-ZNF384* fusion gene in the patient ALL_613. D) Ideogram of the structure of the translocation between chromosome 7 and 9 in the patient ALL_707 resulting in the *PAX5-ELN* fusion gene, which is shown besides the derived chromosome 9 with the direction of the transcription indicated by an arrow. The chromosomes are drawn to scale using the CyDAS software. (E-F) Validation of the chromosome 7q deletion and derived chromosome 9 by FISH in the patient ALL_707.

The most common fusion gene partners of *ZNF384* are the *TCF3* and *EP300* genes. Linked-read WGS determined the chromosomal breakpoints at base-pair resolution for the balanced translocations t(12;19)(p13.31;p13.3)[TCF3-ZNF384] in ALL_604 and t(12;22)(p13.31;q13.2)[EP300-ZNF384] in ALL_613 (**Figure 5B-C**). The heterozygous deletions expected from karyotyping were resolved by the linked-read WGS data to include most of the q-arm of chromosome 7 (7q21.3-q36.3) and chromosome 6q16.2-q22.33 in ALL_604. Although both Long Ranger and CNVnator failed to call the two aforementioned deletions, they were visible in both the linked-read sequencing coverage and Infinium array data (**Supplementary Figure S9**). In ALL_613, a heterozygous deletion of chromosome 16q21-q24.3 and an amplification of the entire q arm of chromosome 1 were observed in the linked-read data. No other large-scale aberrations were detected in the two *ZNF384*-rearranged cases.

One patient with a *PAX5-ELN* fusion gene (ALL_707) detected by RNA-sequencing and short-read WGS was included (11). The karyotype indicated two derived chromosomes (chromosome 7 and 9) as well as a 9p deletion. Using linked-read WGS, we were able to better resolve the aberrations including the translocation t(7;9)(q11;p13) resulting in a derived chromosome 9 harboring the *PAX5-ELN* fusion gene, a truncated chromosome 7, as well as a heterozygous deletion of the chromosome 9p arm with the breakpoint in the *PAX5* locus on 9p13 (**Supplementary Figure S10**). The structure of the resulting derived chromosomes based on the linked-read WGS data is outlined together with FISH validation in **Figure 5D-F**.

### T-ALL

Based on karyotype, a bi-allelic deletion of chromosome 9p21 and two translocations were expected involving chromosomes 7 and 9 and chromosomes 7 and 11 in ALL_559. The homozygous deletion of chromosome 9p21 was clearly resolved in the linked-read WGS data (**Supplementary Figure S11**). Previously generated short-read WGS and RNA-sequencing data identified two translocations involving the T-cell receptor beta locus (*TRBC2* gene) on chromosome 7, namely t(7;11)(q34;p15)[RIC3-TRBC2] and t(7;9)(q34;q31) resulting in the fusion of *TRBC2* to a non-annotated transcript expressed on chromosome 9 between the *TAL2* and *TMEM38B* genes (11). The linked-read data clarified that the two different alleles of *TRBC2* were involved in independent translocation events, as opposed to two different parts of the same allele in the case of a reciprocal event. First, the t(7;11)(q34;p15) resulting in expression of *RIC3-TRBC2* was a consequence of a balanced translocation of chromosome 7 involving one allele of *TRBC2* (**Figure 6A**). On the other allele of *TRBC2*, the t(7;9)(q34;q31) was accompanied by a 0.2 Mb deletion flanked by an inversion of chromosome 7q34 (**Figure 6B-C**), a re-arrangement that was missed by both karyotyping and short-read WGS (11). FISH verified the derived chromosomes determined by linked-read WGS (**Figure 6D-F**).

**Figure 6.**
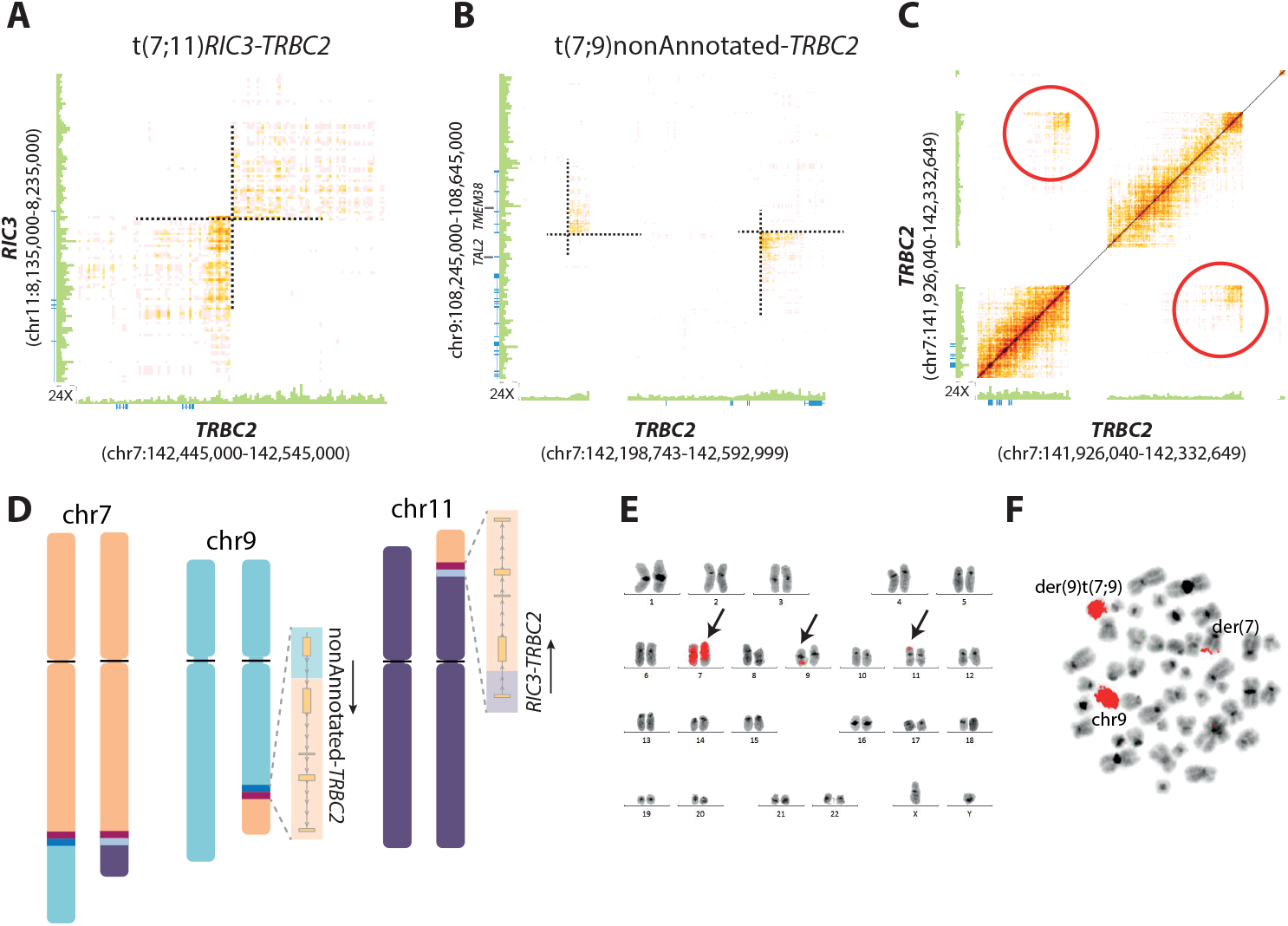
Chromosomal aberrations in the patient ALL_559 (T-ALL) determined by linked-read WGS. (A-C) Heatmaps from the 10x Genomics Loupe software of overlapping linked-reads indicating genomic rearrangements. (A) A balanced translocation between chromosomes 7 and 11. (B) A translocation between chromosomes 7 and 9, which is accompanied by a 0.2 Mb deletion flanked by an inversion of chromosome 7q34 on the second allele at the *TRBC2* locus. The translocation results in an expressed fusion gene between *TRBC2* and a non-annotated gene located 500 bp upstream of *TMEM38B* on chromosome 9. (C) Zoomed in view of the inversion flanking the *TRBC2* locus on 7q34. (D) Ideogram of the structure of the translocations observed in ALL_559. The chromosomes are drawn to scale using the CyDAS software. (E) Whole chromosomal paint depicting the translocation of material from chromosome 7 to chromosomes 9 and 11. (F) Whole chromosomal paint of chromosome 9 depicting the balanced translocation involving chromosome 7.

### Detection of key diagnostic deletions for ALL

To further demonstrate that linked-read WGS is useful for detecting other types of aberrations than large-scale aneuploidies and translocations, we screened the 13 genomes (18 sequencing libraries) for focal deletions in a set of relevant genes for ALL including *BTG1, CDKN2A/B, EBF1, ETV6, IKZF1, PAX5, RB1 and ERG* (35) (**Supplementary Figure S12**). Deletions of each gene were observed in at least one patient, with the exception of *RB1*. All of the deletions identified by linked-read WGS were verified by Infinium arrays. In the paired linked-read WGS libraries generated in different DNA samples from the same patient (REH.1, REH.2 and ALL_402.1, ALL_402.2) or created with different linked-read library preparation protocols (ALL_47.1, ALL_47.2 and ALL_390.1, ALL_390.2) identical deletion patterns were observed. The only exception was the compound heterozygous *ERG* deletion in ALL_390 that was detected in the Chromium library ALL_390.2, but not in the GemCode library ALL_390.1, presumptively due to the lower sequence coverage. The three t(12;21) cases harbored *ETV6* deletions on the allele that was not affected by the translocation, thus resulting in bi-allelic disruption of *ETV6* in all cases. Consistent with previous studies (32,36), recurrent *BTG1* and *IKZF1* deletions were detected in the t(12;21) and *DUX4-IGH* patients, respectively (**Supplementary Figure S13**).

## DISCUSSION

Herein we present the first ALL genomes to be analyzed by “linked-read” whole-genome sequencing technology. Linked-reads enabled highly accurate resolution of the majority of the genomic aberrations defined by cytogenetic methods and refined or identified new structural rearrangements in the 13 analyzed ALL genomes. Although the ALL subtypes and numbers of samples included in the present study are modest, these results provide a strong proof of principle for linked-read WGS for digital karyotyping in ALL. Linked-read WGS is likely to be applicable to other types of malignancies where translocations and aneuploidies are common. Studies taking a similar approach in other cancer types such as triple negative breast cancer (37), metastatic gastric tumors (38) and cell lines (6) have reached similar conclusions.

Linked-read WGS requires long input DNA molecules to gain the most benefit from the technology (3). However, when working with clinical samples, high molecular weight DNA extraction and handling of HMW DNA is not practical in most clinical settings. Therefore, we chose to utilize DNA from patient samples that were prepared using a commonly used column-based DNA extraction method. Although the average length of the DNA was lower than the 50 Kb recommendation by the vendor 10x Genomics, our results show that DNA samples of suboptimal quality are also highly informative for detection genomic aberrations with linked-read WGS, with the exception of whole-genome amplified (WGA) DNA. In all instances where we compared HMW DNA to DNA from standard column extractions, and in most cases where we compared low-coverage GemCode to Chromium library preparation, the results were concordant. Although HMW DNA enabled phasing over chromosomal breakpoints, which makes interpretation of the chromosomal structure and organization easier, long DNA molecules and high sequencing depth may not be required for accurate detection of prognostically relevant aberrations present in the major clone of leukemic samples. In a linked-read WGS study on 23 cell lines, copy number variants were still detectable after sequencing down-sampling to as little as 1-2x coverage, whilst balanced events required approximately 10x coverage (6). It should be noted that these results were derived from HMW DNA extracted from cell lines, and that the results from HMW DNA from cell lines may not be applicable to real clinical specimens with shorter DNA fragments and varying percentages of contaminating normal cells.

In the present study, we focused on detecting large-scale structural aberrations, which are the most relevant type of aberrations for clinical care in ALL (39). We did not address rare somatic SNVs or small insertion-deletions (indels) that are also detectable in linked-read WGS data, which may be of clinical relevance in ALL (11,12,40). Based on our results, we believe that improvements of WGS library preparation and analysis introduced by linked-read technology, has the potential to capture the total load of all types of genomic variants in a single test. Although this new technology is in its infancy, we expect that digital karyotyping by WGS will replace, or at the least complement traditional clinical diagnostic methods such as G-banding and FISH in the future.

## Availability of data and materials

The linked-read WGS, RNA-sequencing and 450k DNA methylation array data sets from the REH cells have been deposited at GEO (GSE116057, submission in progress). Previously generated data from Infinium HumanMethylation450 BeadChips (450k arrays) are available at the GEO (GSE49031). The copy number data generated from the 450k DNA methylation array and SNP array used to determine copy number alterations (CNA) in the ALL patients is available at GEO (submission in progress). The patient/parent consent does not cover depositing data that may be used for large-scale determination of germline variants in a repository. The ALL samples were collected 10-20 years ago from pediatric patients aged 2-15 years, some whom have deceased. The linked-read WGS data and other dataset analyzed in the study are available upon reasonable request from the corresponding author. A summary of additional data sets available for each patient is provided in Supplementary Table S1.

## Authors’ contributions

JN and ACS designed the study. JN, YMZ, and AL analyzed data. AR and TM performed experiments. JA, UNN, and GL provided clinical material and karyotyping data. LC performed FISH experiments and provided expertise on karyotyping. JN, YMZ, and ACS wrote the paper.

## Acknowledgements

Sequencing and SNP genotyping was performed by the SNP&SEQ Technology Platform, which is part of Science for Life Laboratory and the National Genomics Infrastructure at Uppsala University, supported by the Swedish Research Council (VR-RFI) and the Knut and Alice Wallenberg Foundation. Computational analysis was performed using resources provided by SNIC Uppsala Multidisciplinary Center for Advanced Computational Science. We especially thank our colleagues from NOPHO and the ALL patients who contributed samples to this study.

This work was supported by grants from the Selanders Stiftelse (JN), the Swedish Cancer Society (CAN2013/504, CAN2016/559; ACS), the Swedish Childhood Cancer Foundation (PR2014-0100; ACS), and the Swedish Research Council for Science and Technology (E0226301; ACS).

## REFERENCES

1. Sheikine Y, Kuo FC, Lindeman NI. Clinical and Technical Aspects of Genomic Diagnostics for Precision Oncology. Journal of clinical oncology: official journal of the American Society of Clinical Oncology 2017;35:929–33

2. Porubsky D, Garg S, Sanders AD, Korbel JO, Guryev V, Lansdorp PM, et al. Dense and accurate whole-chromosome haplotyping of individual genomes. Nature communications 2017;8:1293

3. Zheng GX, Lau BT, Schnall-Levin M, Jarosz M, Bell JM, Hindson CM, et al. Haplotyping germline and cancer genomes with high-throughput linked-read sequencing. Nature biotechnology 2016;34:303–11

4. Weisenfeld NI, Kumar V, Shah P, Church DM, Jaffe DB. Direct determination of diploid genome sequences. Genome research 2017;27:757–67

5. Mostovoy Y, Levy-Sakin M, Lam J, Lam ET, Hastie AR, Marks P, et al. A hybrid approach for de novo human genome sequence assembly and phasing. Nature methods 2016;13:587–90

6. Garcia S, Williams S, Herschleb J, Marks P, Xu AW, Schnall-Levin M, et al. Linked-Read Sequencing for Molecular Cytogenetics. J Mol Diagn 2017;19:945–

7. Iacobucci I, Mullighan CG. Genetic Basis of Acute Lymphoblastic Leukemia. Journal of Clinical Oncology 2017;0:JCO.2016.70.7836

8. Schmiegelow K, Forestier E, Hellebostad M, Heyman M, Kristinsson J, Soderhall S, et al. Long-term results of NOPHO ALL-92 and ALL-2000 studies of childhood acute lymphoblastic leukemia. Leukemia 2010;24:345–54

9. Moorman AV. The clinical relevance of chromosomal and genomic abnormalities in B-cell precursor acute lymphoblastic leukaemia. Blood reviews 2012;26:123–35

10. Pui CH, Yang JJ, Hunger SP, Pieters R, Schrappe M, Biondi A, et al. Childhood Acute Lymphoblastic Leukemia: Progress Through Collaboration. Journal of Clinical Oncology 2015;33:2938–U24

11. Lindqvist CM, Nordlund J, Ekman D, Johansson A, Moghadam BT, Raine A, et al. The mutational landscape in pediatric acute lymphoblastic leukemia deciphered by whole genome sequencing. Human mutation 2015;36:118–28

12. Holmfeldt L, Wei L, Diaz-Flores E, Walsh M, Zhang J, Ding L, et al. The genomic landscape of hypodiploid acute lymphoblastic leukemia. Nature genetics 2013;45:242–52

13. Tran AN, Taylan F, Zachariadis V, Ofverholm II, Lindstrand A, Vezzi F, et al. High-resolution detection of chromosomal rearrangements in leukemias through mate pair whole genome sequencing. Plos One 2018;13

14. Gianfelici V, Chiaretti S, Demeyer S, Di Giacomo F, Messina M, La Starza R, et al. RNA sequencing unravels the genetics of refractory/relapsed T-cell acute lymphoblastic leukemia. Prognostic and therapeutic implications. Haematologica 2016;101:941–50

15. Lilljebjorn H, Henningsson R, Hyrenius-Wittsten A, Olsson L, Orsmark-Pietras C, von Palffy S, et al. Identification of ETV6-RUNX1-like and DUX4-rearranged subtypes in paediatric B-cell precursor acute lymphoblastic leukaemia. Nature communications 2016;7:11790

16. Yasuda T, Tsuzuki S, Kawazu M, Hayakawa F, Kojima S, Ueno T, et al. Recurrent DUX4 fusions in B cell acute lymphoblastic leukemia of adolescents and young adults. Nature genetics 2016;48:569–74

17. Marincevic-Zuniga Y, Zachariadis V, Cavelier L, Castor A, Barbany G, Forestier E, et al. PAX5-ESRRB is a recurrent fusion gene in B-cell precursor pediatric acute lymphoblastic leukemia. Haematologica 2016;101:e20–3

18. Nordlund J, Backlin CL, Zachariadis V, Cavelier L, Dahlberg J, Ofverholm I, et al. DNA methylation-based subtype prediction for pediatric acute lymphoblastic leukemia. Clinical epigenetics 2015;7:11

19. Marincevic-Zuniga Y, Dahlberg J, Nilsson S, Raine A, Nystedt S, Lindqvist CM, et al. Transcriptome sequencing in pediatric acute lymphoblastic leukemia identifies fusion genes associated with distinct DNA methylation profiles. Journal of hematology & oncology 2017;10:148

20. Biondi A, Schrappe M, De Lorenzo P, Castor A, Lucchini G, Gandemer V, et al. Imatinib after induction for treatment of children and adolescents with Philadelphia-chromosome-positive acute lymphoblastic leukaemia (EsPhALL): a randomised, open-label, intergroup study. The Lancet Oncology 2012;13:936–45

21. Rosenfeld C, Goutner A, Choquet C, Venuat AM, Kayibanda B, Pico JL, et al. Phenotypic characterisation of a unique non-T, non-B acute lymphoblastic leukaemia cell line. Nature 1977;267:841–3

22. Milani L, Lundmark A, Kiialainen A, Nordlund J, Flaegstad T, Forestier E, et al. DNA methylation for subtype classification and prediction of treatment outcome in patients with childhood acute lymphoblastic leukemia. Blood 2010;115:1214–25

23. Paulsson K, Johansson B. High hyperdiploid childhood acute lymphoblastic leukemia. Genes, chromosomes & cancer 2009;48:637–60

24. Abyzov A, Urban AE, Snyder M, Gerstein M. CNVnator: an approach to discover, genotype, and characterize typical and atypical CNVs from family and population genome sequencing. Genome research 2011;21:974–84

25. Obenchain V, Lawrence M, Carey V, Gogarten S, Shannon P, Morgan M. VariantAnnotation: a Bioconductor package for exploration and annotation of genetic variants. Bioinformatics 2014;30:2076–8

26. Hiller B, Bradtke J, Balz H, Rieder H. CyDAS: a cytogenetic data analysis system. Bioinformatics 2005;21:1282–3

27. Nicorici DS, M.; Edgren, H.; Kangaspeska, S.; Murumagi, A.; Kallioniemi, O.; Virtanen, S.;Kilkku, O. FusionCatcher - a tool for finding somatic fusion genes in paired-end RNA-sequencing data. bioRxiv doi: http://dxdoiorg/101101/011650 2014

28. Nordlund J, Backlin CL, Wahlberg P, Busche S, Berglund EC, Eloranta ML, et al. Genome-wide signatures of differential DNA methylation in pediatric acute lymphoblastic leukemia. Genome biology 2013;14:r105

29. Marzouka NA, Nordlund J, Backlin CL, Lonnerholm G, Syvanen AC, Carlsson Almlof J. CopyNumber450kCancer: baseline correction for accurate copy number calling from the 450k methylation array. Bioinformatics 2016;32:1080–2

30. Rasmussen M, Sundstrom M, Goransson Kultima H, Botling J, Micke P, Birgisson H, et al. Allele-specific copy number analysis of tumor samples with aneuploidy and tumor heterogeneity. Genome biology 2011;12:R108

31. Liu YF, Wang BY, Zhang WN, Huang JY, Li BS, Zhang M, et al. Genomic Profiling of Adult and Pediatric B-cell Acute Lymphoblastic Leukemia. EBioMedicine 2016;8:173–83

32. Zhang J, McCastlain K, Yoshihara H, Xu B, Chang Y, Churchman ML, et al. Deregulation of DUX4 and ERG in acute lymphoblastic leukemia. Nature genetics 2016;48:1481–9

33. Clappier E, Auclerc MF, Rapion J, Bakkus M, Caye A, Khemiri A, et al. An intragenic ERG deletion is a marker of an oncogenic subtype of B-cell precursor acute lymphoblastic leukemia with a favorable outcome despite frequent IKZF1 deletions. Leukemia 2014;28:70–7

34. Harvey RC, Mullighan CG, Wang X, Dobbin KK, Davidson GS, Bedrick EJ, et al. Identification of novel cluster groups in pediatric high-risk B-precursor acute lymphoblastic leukemia with gene expression profiling: correlation with genome-wide DNA copy number alterations, clinical characteristics, and outcome. Blood 2010;116:4874–84

35. Moorman AV, Enshaei A, Schwab C, Wade R, Chilton L, Elliott A, et al. A novel integrated cytogenetic and genomic classification refines risk stratification in pediatric acute lymphoblastic leukemia. Blood 2014;124:1434–44

36. Schwab CJ, Chilton L, Morrison H, Jones L, Al-Shehhi H, Erhorn A, et al. Genes commonly deleted in childhood B-cell precursor acute lymphoblastic leukemia: association with cytogenetics and clinical features. Haematologica 2013;98:1081–8

37. Kawazu M, Kojima S, Ueno T, Totoki Y, Nakamura H, Kunita A, et al. Integrative analysis of genomic alterations in triple-negative breast cancer in association with homologous recombination deficiency. PLoS genetics 2017;13:e1006853

38. Greer SU, Nadauld LD, Lau BT, Chen J, Wood-Bouwens C, Ford JM, et al. Linked read sequencing resolves complex genomic rearrangements in gastric cancer metastases. Genome medicine 2017;9:57

39. Janeway KA, Place AE, Kieran MW, Harris MH. Future of clinical genomics in pediatric oncology. Journal of clinical oncology: official journal of the American Society of Clinical Oncology 2013;31:1893–903

40. Grobner SN, Worst BC, Weischenfeldt J, Buchhalter I, Kleinheinz K, Rudneva VA, et al. The landscape of genomic alterations across childhood cancers. Nature 2018;555:321–7

